# Interpretable Antibody–Antigen Structural Interface Prediction via Adaptive Graph Learning and Cyclic Transfer

**DOI:** 10.64898/2026.04.09.717547

**Authors:** Xing Liu, Jason Kantorow, Ashesh K. Chattopadhyay, Srirupa Chakraborty

## Abstract

Experimental structural methods can identify antibody–antigen interfaces with high precision, but they remain time-consuming and resource-intensive, limiting their application across the rapidly expanding space of antibody and antigen sequences. Computational models capable of predicting these interfaces could therefore accelerate antibody discovery and provide insight into the principles governing immune recognition. However, this problem remains challenging due to limited structural datasets, severe class imbalance, and the complex, non-local nature of biomolecular interactions. Here we present VASCIF (Variable-domain Antibody–antigen Structural Complex Interface Finder), a structure-aware framework built on a Masked Graph Attention (MGA) architecture that represents protein complexes as residue graphs and captures long-range structural dependencies through attention-based message passing. The framework is straightforward to implement and enables efficient inference, allowing substantially faster predictions than other existing structure-based approaches. Evaluated on curated structural complexes across multiple benchmark datasets using rigorous cross-validation, VASCIF achieves state-of-the-art performance for residue-level interface prediction. Interpretability analyses reveal that the model recovers biophysically meaningful interaction patterns consistent with known principles of antibody recognition, and redefining interfaces using larger residue distance thresholds (∼10 Å) significantly improves predictive performance. Together, VASCIF provides a practical predictive framework and new insights into antibody–antigen molecular recognition.

## INTRODUCTION

Understanding how antibodies recognize antigens is a central problem in molecular and computational biology, with implications for therapeutic antibody engineering, vaccine design, and immunodiagnostics. Antibodies (Ab), or immunoglobulins, are Y-shaped glycoproteins produced by B cells that play a central role in adaptive immunity^1, 2^. They recognize molecular targets derived from pathogens or host cells and mediate neutralization, opsonization, and immune effector activation^3, 4^. The antigen-binding site of an antibody is formed by the variable regions of its heavy and light chains, which engage a complementary region on the antigen, termed the epitope **(Figure 1)**, through hydrogen bonding, hydrophobic contacts, and electrostatic interactions. The structural principles governing antibody–antigen (Ab–Ag) recognition underpin immune specificity and guide antibody optimization and vaccine design.

**Figure 1:**
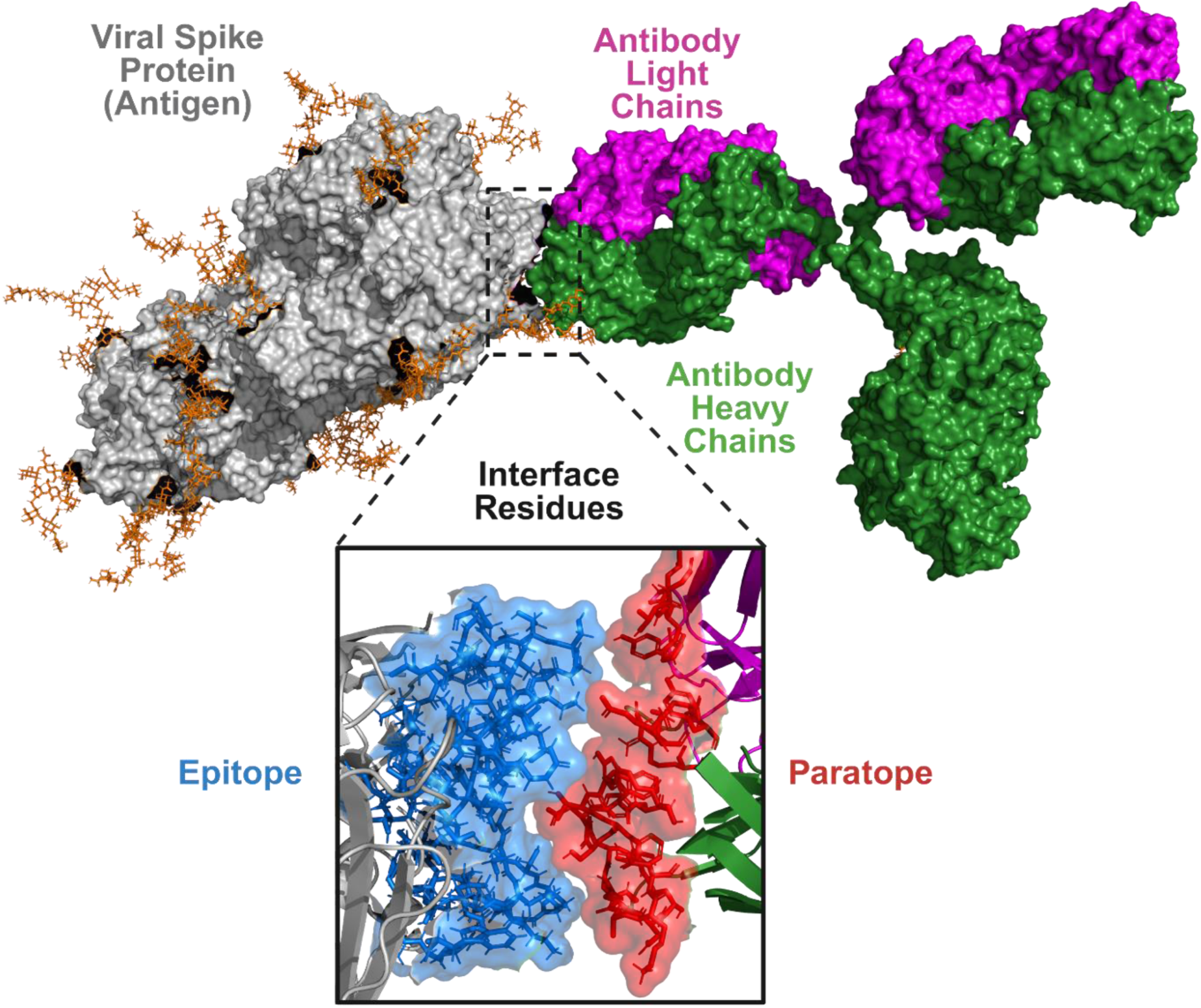
Structural basis of antibody–antigen recognition. Surface representation of an antibody bound to an antigen protein (SARS CoV-2 Spike used as example). The antigen is shown in grey, while the antibody heavy chains are colored green and the light chains magenta. The antibody engages the antigen through residues located within its variable domains. The inset highlights the molecular interface, where antigen residues forming the epitope (blue) interact with the antibody paratope (red) primarily located within the complementarity-determining regions (CDRs) of the heavy and light chains. This interface illustrates the localized yet structurally complex region that mediates antibody–antigen recognition.

Accurately predicting Ab–Ag binding interfaces at residue resolution remains a longstanding and practically important problem. Traditional computational approaches, including molecular docking^5, 6, 7^ and physics-based simulations^8^, provide mechanistic insights but are computationally intensive and often fail to produce unique solutions without initial information about some residues from the interface^5, 6, 7^. The combinatorial diversity of antibodies and viral antigens, the flexibility of complementarity-determining regions (CDRs), and allosteric effects create a high-dimensional recognition landscape^9, 10^. Interface labels are also severely imbalanced: typically fewer than 5–10% of residues participate in binding, making learning sensitive to noise and overfitting^11^. Recent advances in machine learning have transformed protein structure and interaction modeling. Deep architectures leveraging multiple sequence alignments (MSAs), self-attention mechanisms^12^, and graph neural networks (GNNs^13, 14^) have achieved remarkable success in structure prediction and protein–protein interaction tasks^15, 16, 17^. General protein-protein interaction prediction methods end up producing several decoy structures and fail to perform at par for Ab-Ag interfaces^18, 19, 20^. Several models have been developed specifically for paratope and epitope prediction. PECAN introduced a GNN-based framework capturing spatial residue relationships with attention mechanisms for context awareness^21^. EPI-EPMP explicitly modeled the asymmetry between paratope and epitope tasks using convolutional and message-passing strategies^22^. Paragraph employed equivariant GNN layers to predict paratopes from antibody structure alone and was subsequently expanded using a larger curated dataset^23^. ParaSurf incorporated surface geometry, electrostatics, and hybrid convolutional–transformer architectures to enhance binding-site detection^24^. MIPE further leveraged multimodal contrastive learning to fuse sequence and structural information^25^. Despite these advances, key challenges remain: most models either focus on paratopes or epitopes rather than jointly modeling both, depend on dataset-specific heuristics, or treat all residues with equal importance, limiting robustness under extreme class imbalance as would be expected on large antigen surfaces and across antigen families. Most methods also require substantial computational time per prediction, limiting scalability for large-scale screening.

Here we introduce Masked-Graph-Attention (MGA), a new deep learning framework designed to address these challenges in structure-aware residue-level interface prediction. MGA integrates multi-sequence embeddings derived from MSAs, adjacency-based geometric representations of protein structure, and attention mechanisms to jointly model intra- and inter-chain dependencies. Central to the framework is a Dynamic Masking (DyM) module developed by us that adaptively suppresses low-information residues and amplifies contextually important ones in an end-to-end manner. Unlike heuristic down-sampling or CDR-restricted training, DyM learns intrinsic residue-importance distributions directly from data, enabling improved sensitivity in sparse-label environments while remaining architecture-agnostic and broadly deployable. To enhance generalization in data-limited regimes, we introduce Cyclic Transfer with Soft Restart (CTSR), a lightweight training strategy that alternates between interface prediction and related structural tasks, including secondary structure and contact map prediction. By cyclically transferring shared backbone parameters across tasks, CTSR perturbs the optimization landscape in a controlled manner, helping the model escape narrow local minima while incorporating complementary structural priors. Unlike conventional training pipelines^26, 27^, CTSR is cyclic, modular, and compatible with modest dataset sizes typical of structural immunology.

Building on the MGA framework, we develop VASCIF (Variable-domain Antibody–antigen Structural Complex Interface Finder), an interface prediction tool that integrates dynamic masking and cyclic transfer learning to identify paratopes and epitopes across diverse antibody–antigen complexes. We evaluate VASCIF across three benchmark settings, including the Paragraph-expanded dataset^23^, the MIPE dataset^25^, and VASCO^28^ dataset, a previously curated virus-focused dataset spanning diverse antigen families^29^. To ensure robust assessment, we implement cluster-based data splitting to mitigate sequence similarity–driven leakage and report performance across whole Ab–Ag complexes rather than restricting evaluation to predefined CDR regions. VASCIF achieves highly competitive paratope prediction, and state-of-the-art performance in the more difficult task of epitope prediction, with additional gains from CTSR. Interpretability analyses of DyM masks and attention maps further reveal that the model internalizes biophysically meaningful interaction patterns, emphasizing flexible loop regions and chemically favorable residue combinations consistent with known principles of antibody recognition.

Beyond performance improvements, MGA offers practical flexibility. The architecture treats model depth as a tunable hyperparameter, allowing capacity to be matched to dataset scale rather than fixed *a priori*. The framework is fully reproducible, with preprocessing pipelines, training scripts, and evaluation modules publicly available. Owing to its modular design, the method can be extended to incorporate additional molecular features, including glycans involved in immune recognition and viral evasion, and generalized to other macromolecules such as nucleic acids. Together, these results establish VASCIF as a high-performing and interpretable framework for Ab–Ag interface prediction. More broadly, DyM and CTSR provide general strategies for learning from sparse, heterogeneous, and structure-aware biological data, illustrating how adaptive masking and cyclic task transfer can improve optimization and generalization in molecular machine learning.

## RESULTS

### Overview of VASCIF Framework and data

Only a small fraction of antigen residues participate in binding to a particular antibody, as the antibody Fab domain typically engages the antigen through a compact binding footprint of approximately 10 Å radius^30^, resulting in highly localized epitopes within much larger protein surfaces. We therefore developed our Masked Graph Attention (MGA) model as a structure-aware framework for residue-level antibody–antigen interface prediction under extreme class imbalance and structural heterogeneity. (**Figure 2**) MGA operates on residue graphs constructed from three-dimensional complexes, where nodes encode sequence-derived and structural features and edges capture spatial proximity. By stacking graph attention layers, MGA learns contextualized residue embeddings that integrate long-range structural information, allowing interface propensity to be inferred within the global architecture of the complex rather than as an isolated local signal. To address the severe class imbalance inherent to residue-level interface prediction, MGA introduces adaptive masking to down-weight the overwhelming background of non-interface residues during training, enabling the model to concentrate on sparse, information-rich binding determinants. This not only improves learning under highly skewed label distributions, but also yields more interpretable predictions by highlighting residues that drive interface assignment. Complementing this, a cyclic transfer strategy leverages related structural tasks to reinforce shared geometric and physicochemical representations, improving robustness and generalization across diverse Ab–Ag complexes.

**Figure 2:**
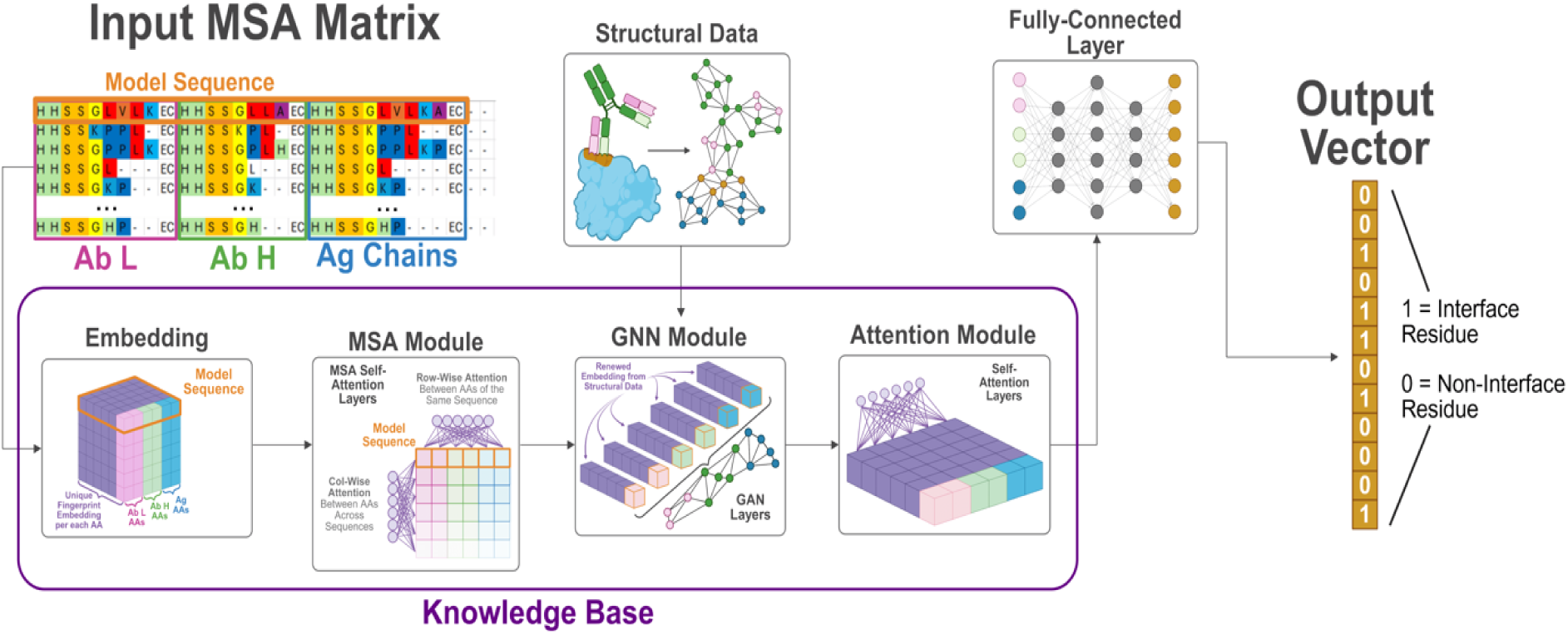
Architecture of the MGA framework for antibody–antigen interface prediction. The model integrates evolutionary and structural information to predict residue-level interface probabilities. Multiple sequence alignments (MSAs) of the antibody light chain (Ab L), antibody heavy chain (Ab H), and antigen (Ag) chains are first encoded into residue embeddings. These representations are processed through an MSA module that captures evolutionary couplings across aligned sequences. Structural information is incorporated through a graph neural network (GNN) module that models residue–residue relationships derived from structural data. The resulting features are further refined by an attention module that learns contextual dependencies across residues. Finally, a fully connected layer produces a binary output vector indicating interface residues (1) and non-interface residues (0). The integrated architecture enables the model to combine sequence conservation, structural connectivity, and contextual attention to identify antibody–antigen interaction sites.

The framework is implemented within VASCIF (Variable-domain Antibody–antigen Structural Complex Interface Finder), which standardizes preprocessing, residue indexing, graph construction, and interface labeling across nonredundant complexes. To allow direct comparison with prior methods, we evaluated VASCIF on the same benchmark datasets, The Paragraph-expanded dataset (1,086 complexes, SAbDab^31^ March 2022 release) was used with merged training–validation sets and 5-fold cross-validation; interface residues comprise ∼6–7% of all residues. The MIPE dataset^25^, curated from SAbDab and filtered to 626 nonredundant complexes using stringent sequence-identity thresholds, follows the standard 90/10 train–test split with 5-fold cross-validation; interface fractions are similarly sparse (∼5–7%). To further test robustness, we additionally used our previously curated virus-focused dataset spanning SARS, MERS, Influenza, Ebola, HIV, and Nipah complexes^28^. To prevent structural leakage, antibody sequences were clustered by similarity and split cluster-wise (80/20). The final curated datasets comprise approximately 854 non-redundant complexes in the training and 202 in the test sets (see **Methods**).

### VASCIF Enables Robust Interface Prediction Under Severe Class Imbalance

Residue-level Ab–Ag interface prediction is intrinsically constrained by extreme class imbalance. Across all benchmark datasets examined here, interface residues constitute only ∼5–10% of total residues, with paratopes typically ∼4–8% and epitopes ∼4–12% depending on the dataset (see Methods). This imbalance reflects the structural nature of antibody recognition: a compact Fab footprint engages a limited antigen surface within a much larger protein scaffold. Naïve classifiers therefore favor majority-class (non-interface) predictions, leading to inflated AUROC values while failing to maintain precision at practical recall thresholds. To address this regime, VASCIF was trained using an imbalance-aware objective combining class-weighted cross-entropy, a differentiable AUPR surrogate, and graph-based smoothness regularization. Because the positive class is sparse, we emphasize AUPR as the primary metric, which better reflects performance under skewed labels^32^.

On the Paragraph-expanded dataset, VASCIF demonstrates consistent improvements in AUPR across successive CTSR training cycles (**Table 1**). For epitope prediction, AUPR increases from 0.472 under baseline training to 0.490 after CTSR. Paratope prediction similarly improves from 0.765 to 0.778. The lower AUPR for antigen residues relative to antibody residues is expected because epitope prediction requires identifying a sparse binding patch across a much larger and more heterogeneous antigen surface, whereas paratopes are confined to the antibody variable domain and thus present stronger intrinsic structural signals. In this setting of extreme class imbalance, even a ∼0.02 increase in AUPR is substantial, as it can move several true interface residues into the top-ranked prediction set while reducing false positives in the regime most relevant for experimental validation. This is particularly important because most current state-of-the-art antibody interface predictors focus primarily on paratope identification, whereas accurate epitope prediction on the antigen remains the more challenging task. The ability of VASCIF to jointly improve prediction on both sides of the interface therefore represents an important practical and conceptual advance. Precision–recall curves show that these gains arise primarily from improved precision at low to moderate recall levels. These levels encompass the regime most relevant for experimental prioritization where minimizing false-positive residues is critical (**Figure 3**). Performance improvement across MIPE and VASCO datasets are also shown in **Table 1**, where the VASCIF model consistently remains the best performing Ab+Ag predictor.

**Table 1:**
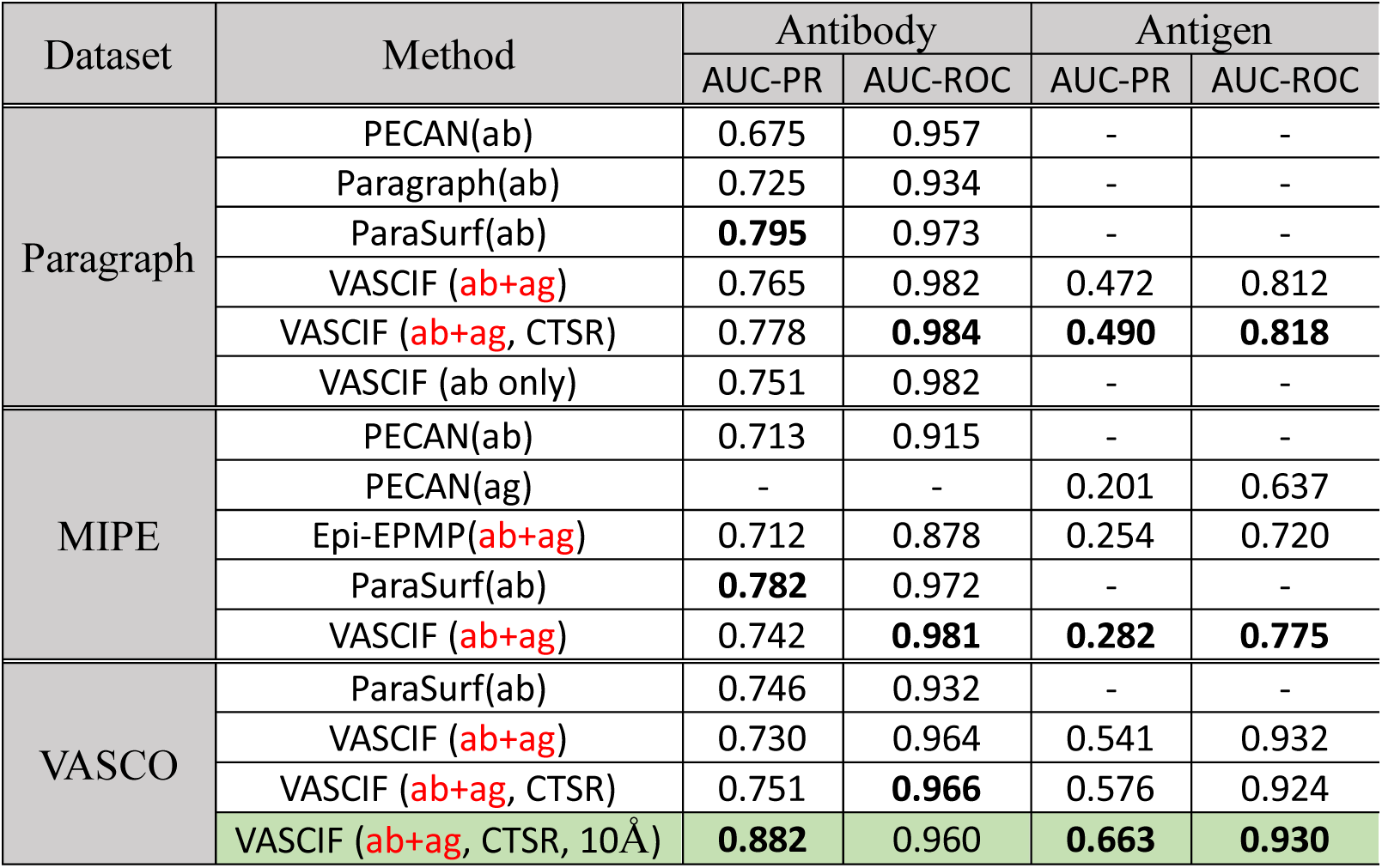
Performance of methods on datasets. Our suggestions for best performing models are shown in bold. Note that most other equivalent performing models only predict Ab residues. The ones that can predict both Ab and Ag are marked in red. The overall best recommended model for predicting both Ab and Ag residues is highlighted in green – the VASIF model with 10 Å search sphere of residues.

**Figure 3:**
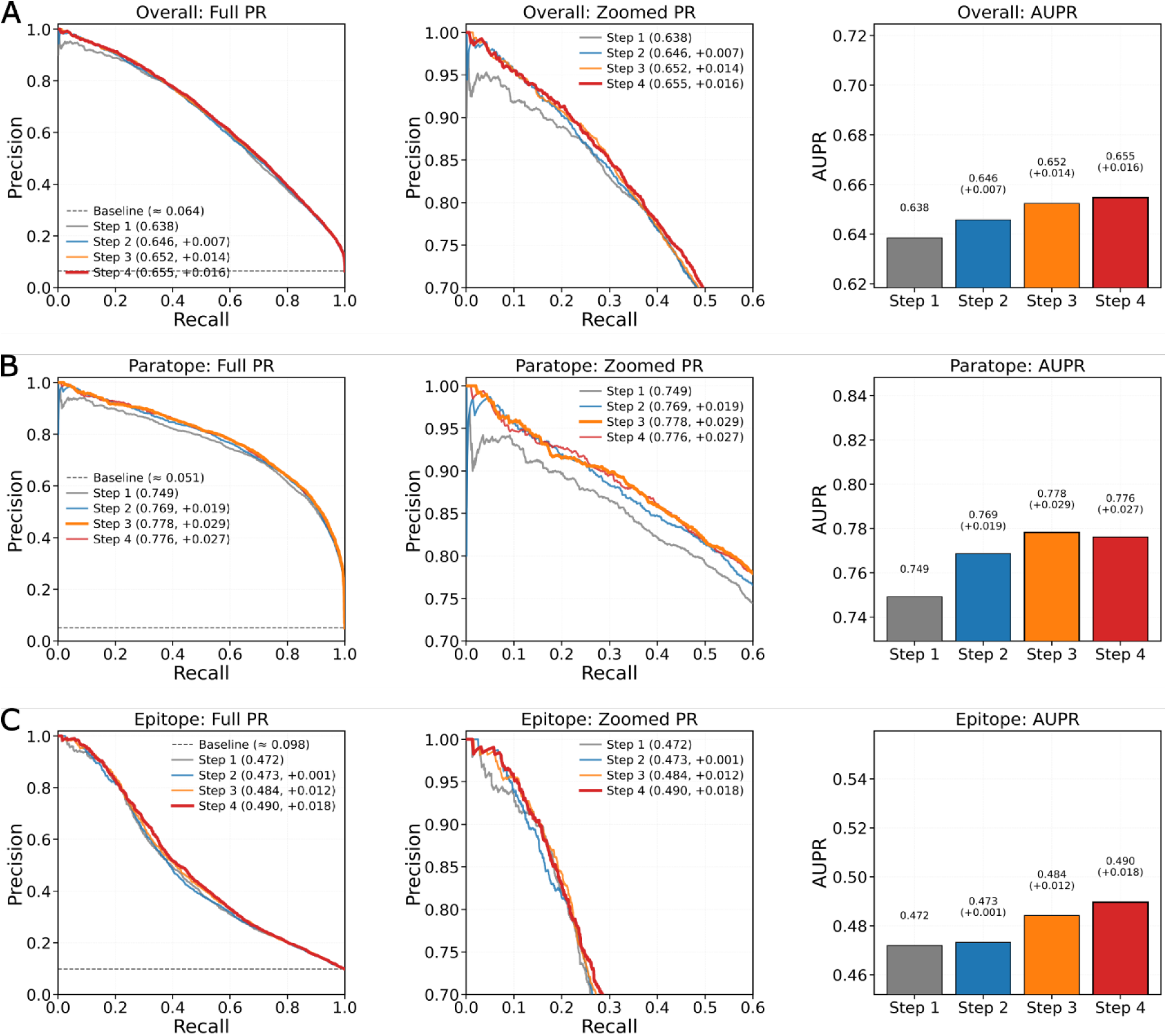
Precision–recall performance across CTSR training steps. Precision–recall (PR) curves and corresponding area under the precision–recall curve (AUPR) for (A) overall residues, (B) paratope residues, and (C) epitope residues are shown, with full PR curves (left), zoomed high-precision regions (middle), and AUPR summaries (right). Step 1 denotes the baseline model, while subsequent steps incorporate iterative refinement. Improvements are consistently observed for epitope prediction (ΔAUPR up to +0.018) and overall performance, indicating enhanced discrimination of antigen-side interface residues. In contrast, paratope performance peaks at Step 3 (ΔAUPR +0.029) and slightly decreases thereafter, reflecting a deliberate reallocation of model capacity toward improving the more challenging epitope prediction task under class imbalance. Dashed lines indicate the baseline positive class fraction.

Ablation analysis clarifies how VASCIF maintains robustness under imbalance (**Figure 3A)**. Removing the GNN module produces the largest degradation in AUPR for both paratopes and epitopes, highlighting the importance of structured message passing in distinguishing true binding clusters from isolated exposed residues. The imbalance-aware design of MGA generalizes beyond a single dataset. On the MIPE dataset, where interface residues represent ∼6% of residues, VASCIF achieves state-of-the-art epitope AUPR (0.282) while maintaining competitive paratope performance. On the VASCO dataset, characterized by heterogeneous viral antigen classes and cluster-based splitting to prevent leakage, MGA preserves strong discrimination even when interface fractions fall to ∼4–6%. Together, these results demonstrate that VASCIF remains robust under severe structural imbalance. By optimizing precision–recall behavior and integrating adaptive masking with graph-aware regularization, VASCIF maintains high precision for sparse binding residues without sacrificing recall.

### Dynamic Masking Improves Sensitivity to Sparse Binding Residues

Training Ab–Ag interface predictors under extreme class imbalance requires mechanisms that emphasize informative structural signals while suppressing background features. To address this challenge, we introduce Dynamic Masking (DyM), a learnable gating mechanism that adaptively rescales residue embeddings during training to emphasize informative positions while suppressing less relevant ones. Despite its simplicity, DyM can be implemented with only a few lines of PyTorch^33^ code (**Figure 4**) and yields substantial performance gains (**Figure 5A**). Mechanistically, DyM applies a sigmoid-activated linear projection to each residue embedding to produce a continuous mask weight that multiplicatively modulates feature representations. Unlike static dropout or heuristic filtering strategies, such as, restricting predictions to complementarity-determining regions (CDRs), DyM is trained end-to-end and adapts to the structural context of each residue, enabling data-driven sparsification of residue representations. Conceptually, DyM differs from several approaches used to handle noisy or imbalanced representations in deep learning. Stochastic regularization methods such as dropout^34^ randomly suppress features during training but do not incorporate structural context or learn which positions are informative. Attention mechanisms^12^ highlight residues through learned weights, yet primarily capture contextual dependencies rather than enforcing sparsity within residue representations. Hard region-filtering strategies used in antibody modeling, such as restricting predictions to CDR loops^23^, rely on predefined structural heuristics and may exclude informative framework residues. In contrast, DyM introduces a learnable gating mechanism related to adaptive sparsification methods^35^, allowing the model to reallocate capacity toward residues contributing to binding discrimination without imposing manual structural constraints.

**Figure 4:**
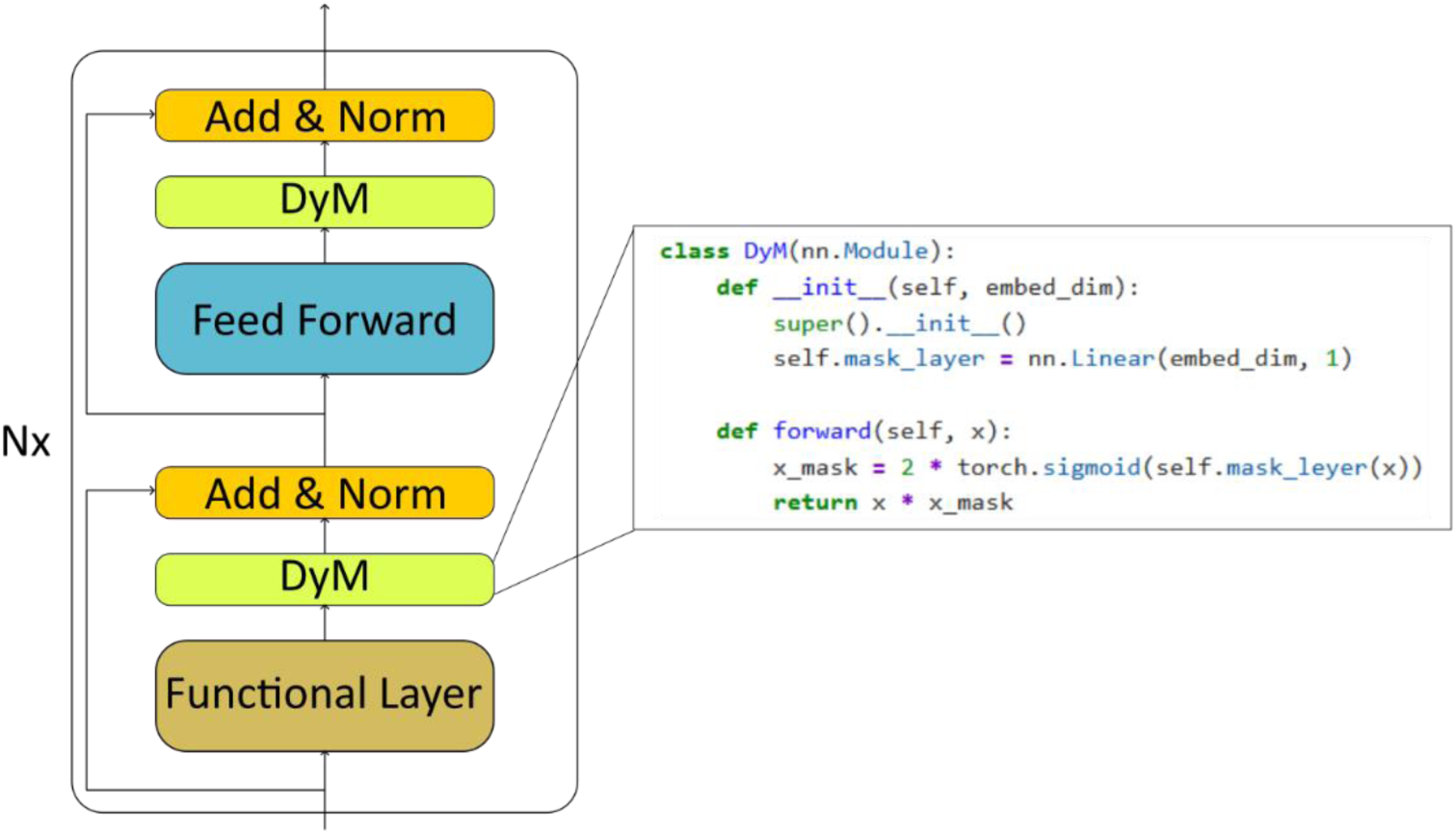
The building block of dynamic masking structures. In addition to the traditional transformer architecture, an additional learnable masking is applied to the functional layer and Feed Froward layer.

**Figure 5:**
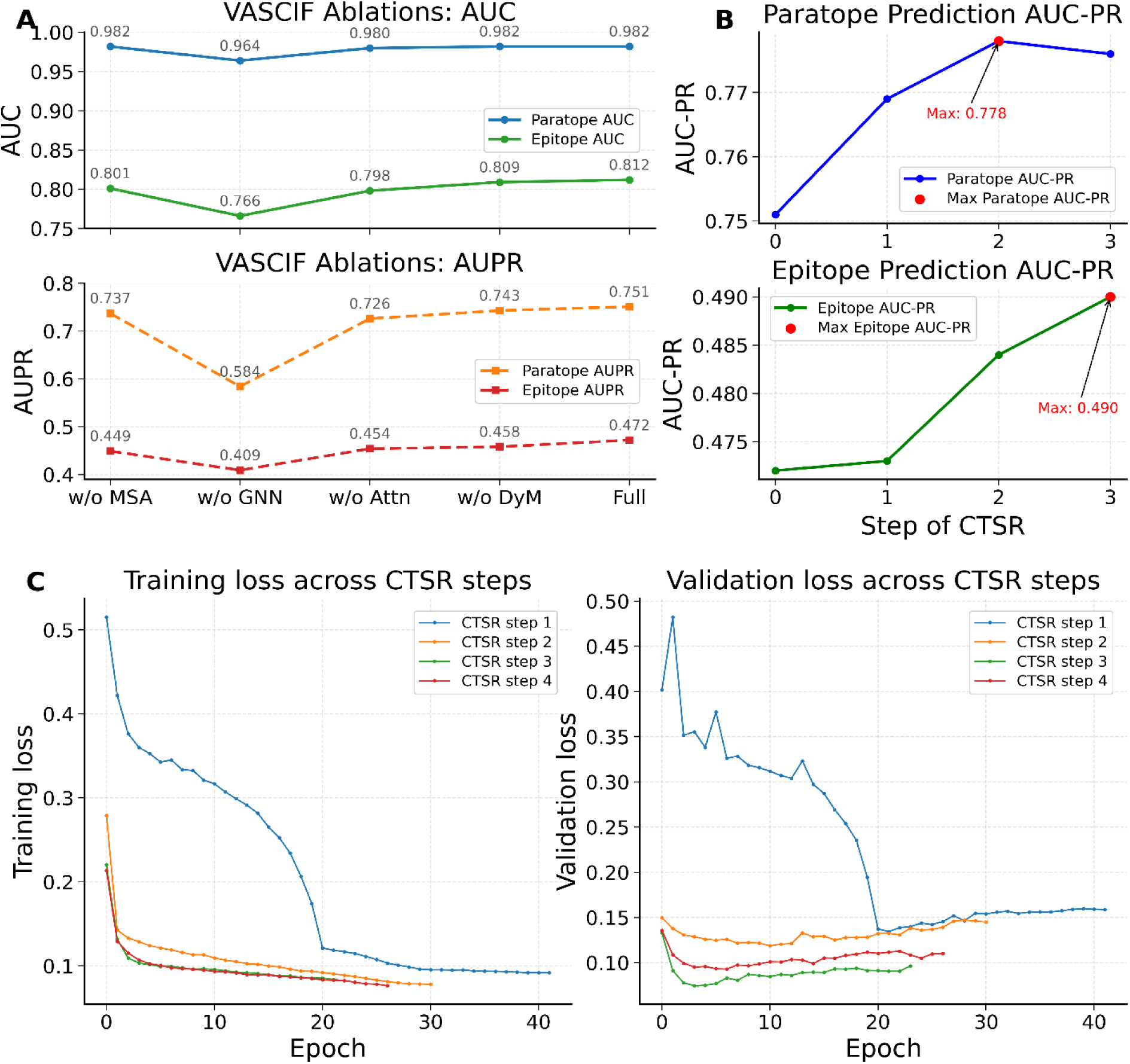
(A) Performances of VASCIF and its variants in joint paratope-epitope prediction in Paragraph dataset. (B) The performance of interface prediction after each CTSR step. (C) Training and validation loss trajectories across CTSR cycles. Each curve corresponds to one stage of cyclic transfer between interface prediction and auxiliary structural tasks. Early stopping based on validation loss was applied to mitigate overfitting.

Ablation analysis confirms the contribution of DyM (**Figure 5A**). Removing DyM reduces epitope AUPR from 0.472 to 0.458 (ΔAUPR = −0.014), one of the largest drops among architectural ablations. Given that only ∼6% of antigen surface residues form the epitopes, a 0.014 drop in AUPR is substantial in practice, reflecting poorer ranking of rare interface residues against a large excess of negatives. In the top-ranked subset typically used for experimental prioritization, this can correspond to several additional false positives and/or missed true epitope residues.

Further analysis of fused mask profiles across layers reveals enrichment of high mask weights in loop regions and depletion in well-packed α-helices and β-sheets. Physically, this indicates that DyM preferentially learnt to retain residues that are solvent-exposed and conformationally adaptable (features that favor intermolecular recognition) while down-weighting residues embedded in rigid, internally stabilized secondary-structure elements that are less likely to contribute directly to binding. These patterns align with known features of Ab–Ag interfaces, where flexible loops and polar or aromatic residues frequently dominate binding interactions.

### Cyclic Transfer Improves Optimization and Generalization

Training deep graph-based models on relatively small structural datasets that comprise only hundreds of complexes, presents a major optimization challenge. Under limited supervision, gradient descent can converge to sharp local minima that overfit dominant structural patterns while failing to generalize to unseen antigen geometries. In practical terms, this means the model can memorize recurrent surface motifs or residue environments seen in the training set, yet fail to recognize alternative geometric arrangements of chemically similar binding determinants on structurally distinct antigens. To address this limitation, we introduce Cyclic Transfer with Soft Restart (CTSR), a training paradigm that alternates between the primary interface prediction task and related structural objectives (**Figure 6**). CTSR first trains the backbone on interface prediction, then transfers learned representations to auxiliary tasks such as secondary structure or contact map prediction. After auxiliary training, the updated backbone is returned to the interface task for further fine-tuning. Training loss trajectories across CTSR cycles (**Figure 5B**) show that initial CTSR transfers substantially reduce validation loss relative to single-task training. Although the final cycle does not further decrease the minimum loss, predictive performance (measured by epitope AUPR) continues to improve, suggesting that CTSR enhances ranking and discrimination of interface residues rather than probability calibration. Empirically, CTSR yields consistent improvements under severe class imbalance. On the Paragraph-expanded dataset, paratope AUPR increases from 0.751 to 0.778 and epitope AUPR from 0.472 to 0.490 without altering model architecture. Conceptually, CTSR differs from conventional multitask learning^36^, where multiple objectives are optimized simultaneously through shared representations. CTSR instead separates tasks temporally, avoiding the need to balance competing gradients. It is also related to cyclic optimization methods such as stochastic gradient descent with warm restarts^37^ and cyclical learning-rate schedules^38^ However, those methods perturb optimization dynamics without changing the learning objective, whereas CTSR introduces complementary supervisory signals that reshape the representation space. Auxiliary tasks follow a structured progression or ‘curriculum^39^’. Training begins with local structural objectives such as secondary structure prediction before progressing to other relational tasks such as residue contact prediction or solvent-accessible structural context characterization, thereby stabilizing training and introducing long-range constraints.

**Figure 6:**
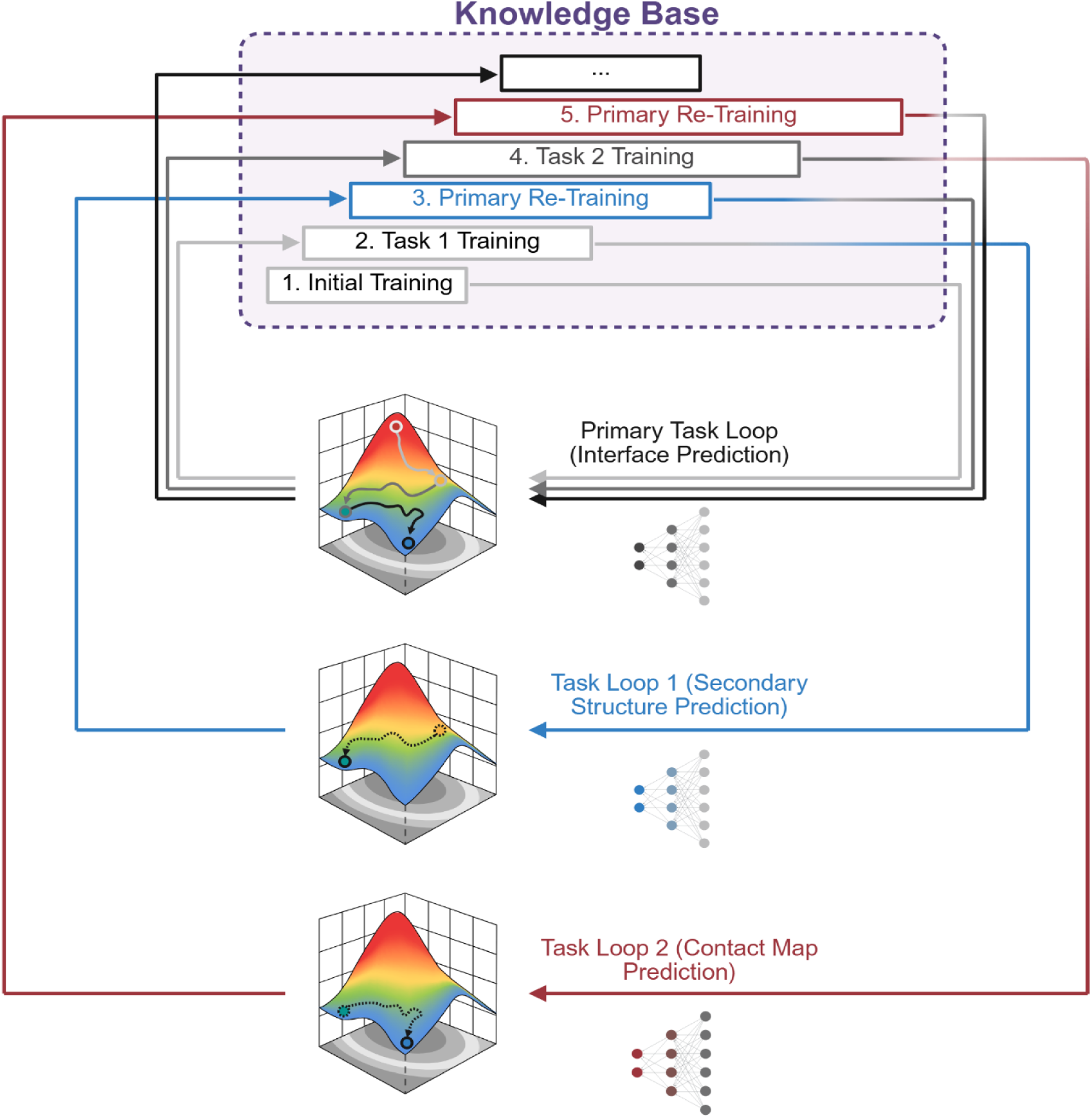
Cyclic Transfer with Soft Restart (CTSR) training framework. Schematic illustration of the CTSR training strategy used to improve interface prediction under limited labeled data.

### Benchmark Performance Across Diverse Antibody–Antigen Datasets

To assess robustness across structural regimes, we benchmarked VASCIF against leading interface prediction methods on three independent datasets: Paragraph-expanded, MIPE, and VASCO. These datasets differ in antigen diversity, splitting strategy, and interface distribution. On the Paragraph-expanded benchmark, VASCIF achieves strong paratope performance and better epitope prediction relative to graph-based and surface-based baselines (**Table 1**). While AUROC values remain high due to dominant negatives, AUPR differentiates models under severe imbalance. Incorporating CTSR further improves performance without altering architecture. On the MIPE dataset, VASCIF attains state-of-the-art epitope AUPR while maintaining competitive paratope performance. Notably, VASCIF predicts both paratopes and epitopes within a unified framework, whereas most prior methods specialize in one interface side. The VASCO dataset provides an additional test of structural heterogeneity. Antibody sequences were clustered and split at the cluster level to prevent sequence leakage. Under this stringent partitioning, VASCIF maintains strong discrimination for both paratopes and epitopes. Across datasets, several patterns emerge: epitope prediction remains the more challenging task due to greater antigen variability; improvements are most pronounced in AUPR rather than AUROC; and performance gains occur without dataset-specific architectural modifications, indicating robust generalization. When trained using a 10 Å interface definition rather than the conventional 4.5 Å cutoff, performance further improves, consistent with both biophysical intuition and reduced label sparsity. Notably, most prior Ab–Ag interface prediction studies have adopted a narrow heavy-atom distance cutoff of ∼4.0–4.5 Å to define interface residues, effectively emphasizing only direct contact geometry. In contrast, when we use a 10 Å definition, it better reflects the physical reality that molecular recognition is governed not only by direct contacts but also by longer-range non-bonded interactions, including electrostatic and van der Waals contributions, which typically extend over larger spatial scales.

### Model scaling demonstrates optimal capacity under data-limited conditions

Modern deep learning systems often show predictable scaling behavior, where larger models improve performance given sufficient data. In contrast, Ab–Ag interface prediction operates in a low-data regime with strong structural redundancy. Naïve scaling can therefore lead to overparameterization and degraded generalization. To examine capacity alignment, we evaluated MGA architectures spanning varying depths of MSA blocks, GNN layers, and attention layers while holding training protocol constant (**Supplementary Table S1**). On the MIPE benchmark, performance shows gradual saturation rather than monotonic improvement. Moderate-depth configurations achieve peak epitope AUPR, whereas deeper models yield marginal or negative returns despite increased parameter count. These results indicate that MGA treats architectural depth as a tunable hyperparameter rather than a fixed design choice, enabling capacity to be aligned with dataset scale.

### Interpretable representations reflect structural determinants of Ab-Ag binding

Beyond predictive accuracy, we examined whether VASCIF learns meaningful biochemical principles governing Ab–Ag recognition through visualization of representative complexes (**Figure 7**). We selected two cases corresponding to the best and worst model predictions we obtained, PDB *8hn6* and *7zlk,* respectively, to assess the robustness of learned representations. The poorer-performing case, 7zlk, is particularly challenging because its interface is distributed across less stereotypical surface features, as in, only the heavy chain of the Fab interacts with the antigen protein, with several viral Env surface glycans also directly participating in Ab-Ag interactions. Across both examples, fused Dynamic Mask (DyM) profiles consistently (**Figures 7C, 7G**) highlight residues in flexible loop regions while suppressing well packed alpha helices and beta sheets, consistent with the established role of complementarity determining region loops in mediating antibody binding^40^. Notably, even in the more challenging case *7zlk*, DyM continues to identify surface loop regions as potential interaction hotspots and masking buried structured motifs despite reduced prediction accuracy. In contrast, last layer attention maps appear diffuse and fail to reliably localize these regions (**Figures 7D, 7H**). Last-layer attention maps are often used as an interpretability proxy in graph-based deep learning models, even though they primarily capture contextual dependencies rather than direct residue-level importance. This contrast suggests that DyM captures intrinsic structural determinants of binding more effectively than attention, and should serve as a useful tool for model explainability, particularly in structural biology settings where identifying physically meaningful interaction regions is critical.

**Figure 7:**
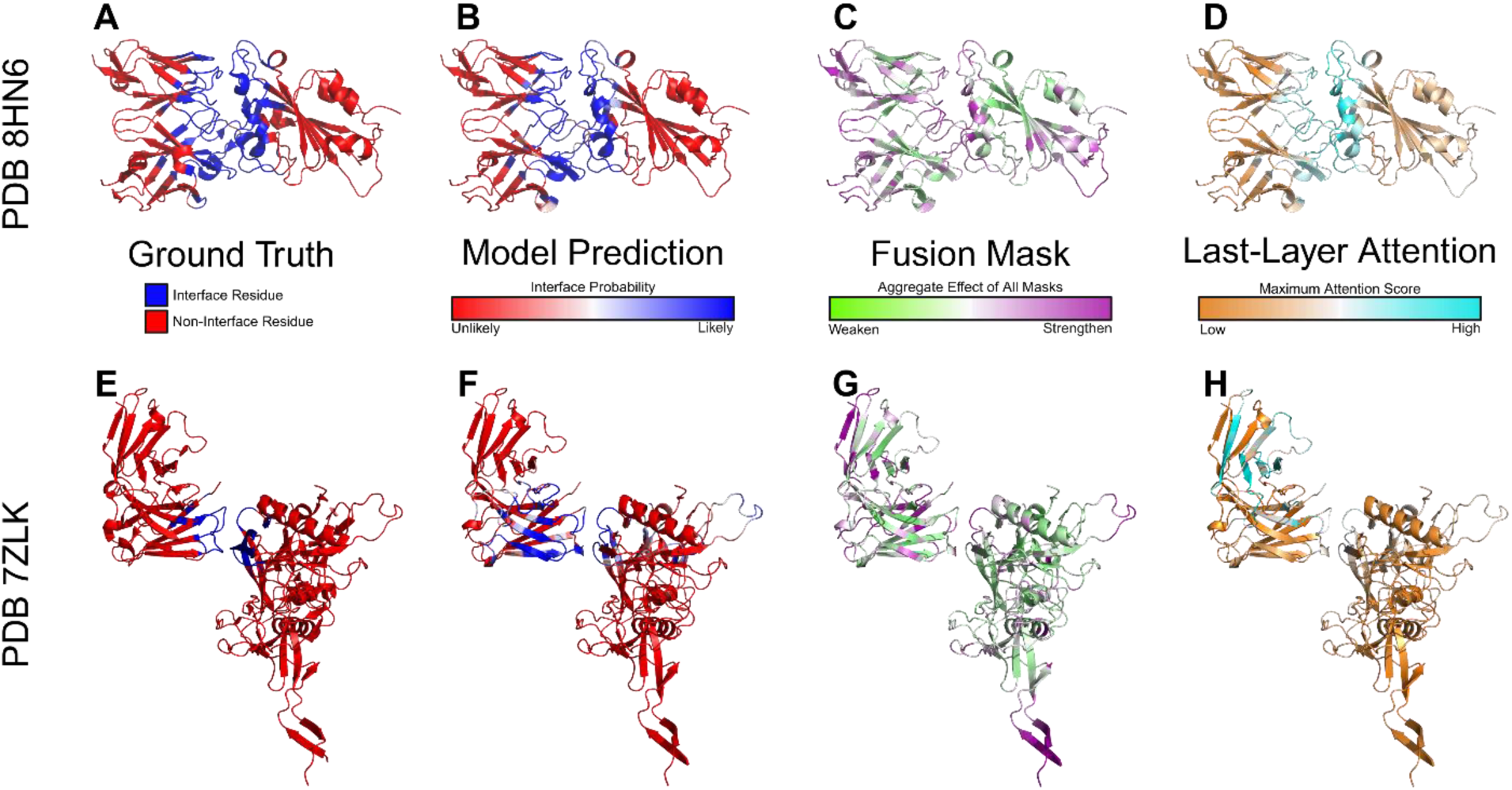
Structural visualization of interface prediction and model interpretability signals. Representative antibody–antigen complexes illustrating ground-truth interfaces, model predictions, and interpretability features learned by VASCIF. Two example complexes are shown. **(A, E)** Ground-truth interface residues, where interface residues are shown in blue and non-interface residues in red. **(B, F)** Model-predicted interface probabilities mapped onto the structures, with blue indicating high interface probability and red indicating low probability. **(C, G)** Fused Dynamic Mask (DyM) profiles showing the aggregate effect of masking across layers, where magenta indicates strengthened residue representations and green indicates suppressed representations. **(D, H)** Residue-level attention scores from the final attention layer, with warmer colors indicating higher attention weights.

Comparison of ground-truth amino acid interaction frequencies with model-predicted interaction propensities (**Figure 8**) shows that the learned interaction matrix recapitulates experimentally observed residue preferences. Tyrosine and serine residues on the antibody side frequently engage diverse antigen residues, while polar residues such as asparagine and glutamine are enriched on the antigen side^41, 42^. These trends are consistent with established principles of antibody–antigen interface chemistry, including aromatic stacking, hydrogen bonding, and polar complementarity. Aromatic residues, especially tyrosine and tryptophan, are known to dominate antibody binding “hot spots” because their side chains provide versatile interaction modes through π–π interactions, hydrogen bonding, and hydrophobic contacts^43, 44^.

**Figure 8:**
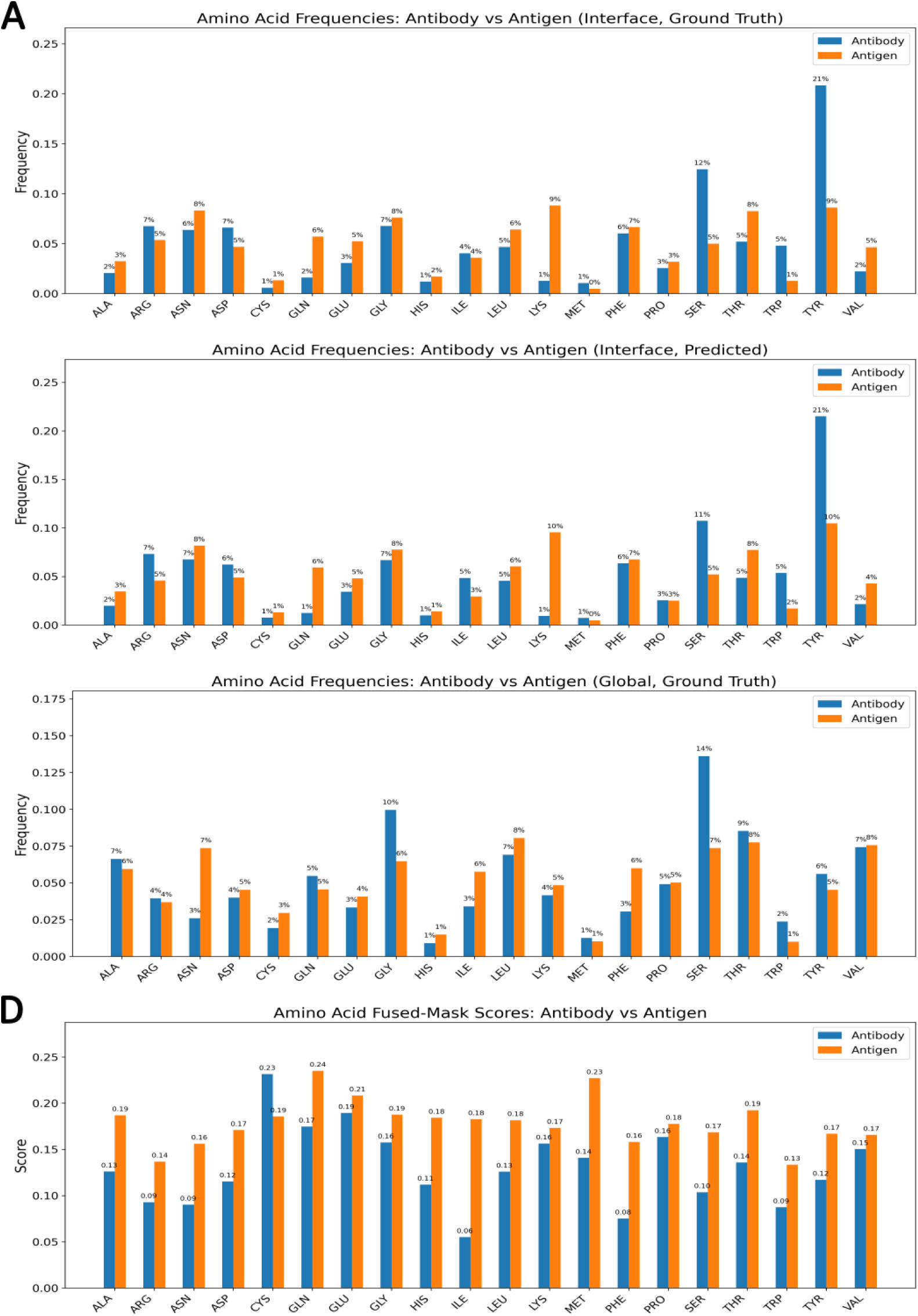
Comparison of Ground-Truth and Predicted Amino Acid Distributions in Antibody–Antigen Complexes. (A) Ground-truth interface composition: normalized frequencies of amino acids observed at antibody–antigen interfaces, where interface residues are defined by a 4.5 Å heavy-atom distance threshold. (B) Predicted interface composition: model-predicted amino acid frequencies at the interface, reflecting the network’s learned residue-level preferences. (C) Global composition: background frequencies of amino acids across all antibody and antigen residues, providing a baseline for enrichment comparison. (D) Fused mask profile: averaged fused-mask values for each amino acid, integrating the contributions from multiple model layers to highlight residues with high predicted importance.

To further examine these trends, we analyzed residue–residue interaction frequency matrices (**Figure *9***). The ground-truth matrix exhibits clear preferences for specific residue interactions, particularly involving aromatic residues such as tyrosine and tryptophan on the antibody side, consistent with their established role in binding hotspots. The model-predicted interaction matrix preserves the overall structure of these interaction patterns, capturing dominant residue preferences while displaying a smoother and more distributed signal. Rather than reproducing sharp, high-intensity peaks, the predicted matrix spreads interaction propensity across neighboring residue types, suggesting that the model captures broader interaction neighborhoods rather than isolated residue-specific contacts. In contrast, the last-layer attention map shows a relatively uniform distribution with limited residue-specific structure, indicating that attention weights do not encode meaningful biochemical interaction preferences at the residue-pair level. These observations suggest that VASCIF learns physically relevant interaction trends while generalizing beyond sparse empirical contact frequencies, and further support that DyM-driven representations, rather than attention alone, encode the structural determinants of antibody–antigen recognition.

**Figure 9:**
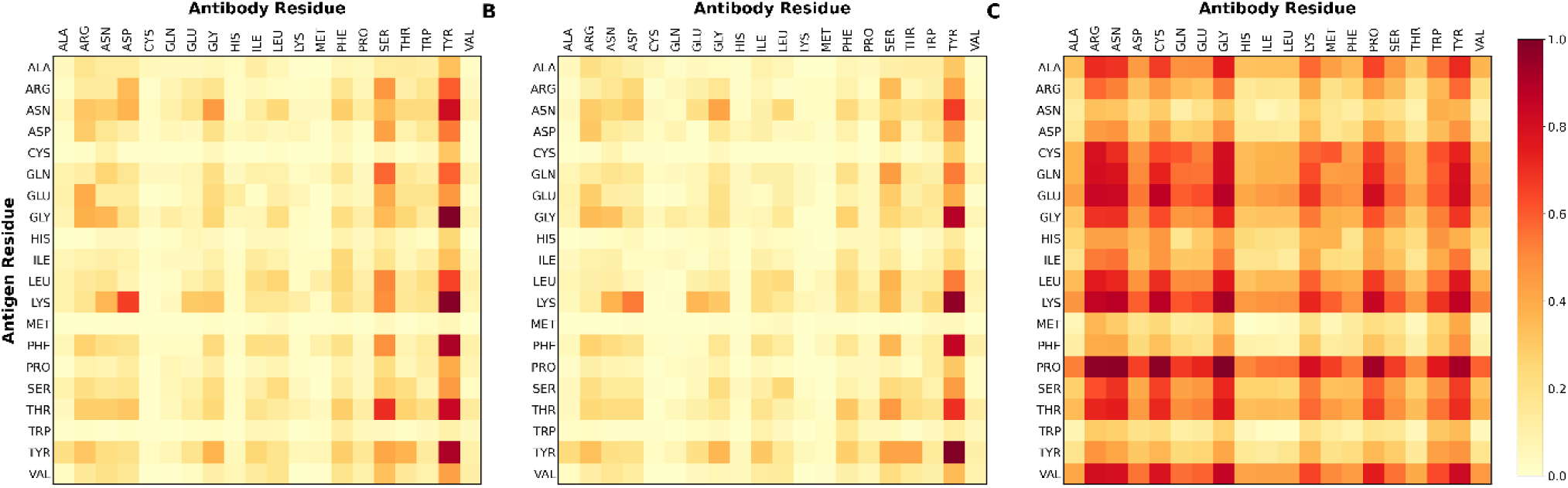
Amino acid interaction patterns learned by VASCIF. Heatmaps comparing ground-truth residue interaction frequencies, model-predicted interaction propensities, and attention-derived interaction patterns for antibody–antigen complexes. The horizontal axis represents antibody residues and the vertical axis represents antigen residues. **(A)** Ground-truth interaction matrix showing normalized frequencies of amino acid pairs observed at antibody–antigen interfaces, where interactions are defined by at least one heavy-atom contact within 4.5 Å. **(B)** Model-predicted interaction matrix showing the average predicted likelihood of residue–residue interactions across antibody–antigen pairs. **(C)** Last-layer attention map showing the average attention weights extracted from the final attention layer of the model. Comparison across panels illustrates that the predicted interaction matrix closely recapitulates experimentally observed amino acid interaction preferences, whereas attention weights capture broader contextual dependencies rather than direct biochemical interaction propensities.

### Antibody-only modeling captures strong intrinsic paratope determinants, but antigen context improves refinement

Recent studies have suggested that accurate paratope prediction may be achievable using antibody information alone, without explicit antigen input^23, 24^. To test this within VASCIF, we compared antibody-only input against full Ab–Ag input using the same architectural depth and training protocol (**Table 1**). Consistent with prior reports, antibody-only modeling already performs strongly (AUPR = 0.751), indicating that antibody variable domains encode substantial intrinsic information about binding propensity. However, including antigen context yields a reproducible improvement (AUPR = 0.765), and performance increases further with CTSR (AUPR = 0.778), showing that partner information provides meaningful additional signal even for paratope prediction. This is biologically intuitive: paratope formation is shaped not only by intrinsic CDR geometry, loop flexibility, and residue chemistry, but also by structural complementarity to the antigen surface. This is consistent with the enrichment of characteristic CDR-associated residue chemistries and loop conformations, including tyrosine- and serine-rich hotspots (**Figure 6**), which predispose specific antibody positions toward interaction even before partner-specific context is considered. The modest absolute gain nevertheless suggests that antibody-only models remain useful for rapid screening when antigen structures are unavailable, whereas full antibody–antigen models are preferable when precise residue-level localization is required. Together, these results support a nuanced view in which intrinsic antibody features dominate paratope signal, but explicit antigen context measurably refines prediction.

## DISCUSSION

Antibody–antigen recognition arises from a localized yet physically complex network of intermolecular interactions involving both short-range contacts and longer-range electrostatic and van der Waals forces. Most prior computational studies define residue-level interfaces using a ∼4.5 Å heavy-atom cutoff, a convention inherited from crystallographic contact definitions. While this threshold captures direct atomic contacts, it represents only a subset of the interactions contributing to binding. In molecular simulations, non-bonded interactions are typically modeled using switching or cutoff distances of ∼10–12 Å to account for electrostatic and dispersion interactions beyond immediate contacts. Consistent with this physical picture, we observe that defining residue-level interaction neighborhoods using a ∼10 Å cutoff improves predictive performance compared with the conventional 4.5 Å definition. This suggests that Ab–Ag interfaces are more appropriately viewed as interaction neighborhoods rather than strict contact sets, and that interface prediction benefits from definitions reflecting the physical range of molecular forces rather than purely geometric contacts. We suggest that future residue-level antibody–antigen interface benchmarks consider moving beyond the conventional 4–4.5 Å standard toward a ∼10 Å interface definition.

More broadly, Ab–Ag interface prediction can be reframed as a sparse structural signal detection problem, in which a small set of functionally important residues must be identified within a large background of structurally similar but non-interacting residues. Antibody binding occurs through a compact footprint localized to a small region of the antigen surface, so informative residues represent only a small fraction of the total structural landscape. The challenge therefore lies in detecting subtle structural signals within overwhelming background. In this context, AUPR values ranging from ∼0.3 to 0.7 may appear modest compared with balanced tasks. However, antibody–antigen interface prediction is intrinsically difficult: across all evaluated datasets only ∼5–7% of residues are labeled as interface residues. Under such extreme class imbalance, metrics such as AUROC can remain deceptively high even for weak classifiers, whereas AUPR provides a more faithful measure of positive-class recovery. In this regime, even modest improvements in AUPR translate into meaningful gains in precision at practical recall thresholds, improving the reliability of candidate interface residues for experimental validation.

DyM (**Figure 4**) addresses this sparsity challenge at the representation level. Rather than restricting predictions to predefined structural regions such as complementarity-determining regions (CDRs) or relying on stochastic negative sampling, DyM introduces a learnable representation-gating mechanism that dynamically reallocates model capacity toward informative residues by adaptively rescaling residue embeddings during training. The resulting fused mask profiles consistently enrich loop regions while suppressing well-packed α-helical and β-sheet environments. These patterns mirror known structural features of antibody–antigen interfaces, where flexible loops frequently mediate binding interactions. Importantly, these preferences emerge without explicit structural priors, suggesting that DyM enables intrinsic importance learning that allocates representational capacity toward residues with consistent binding relevance.

Complementing this architectural component, CTSR addresses optimization challenges associated with limited structural datasets. With datasets comprising only hundreds to low thousands of complexes, conventional training strategies can converge prematurely to sharp minima dominated by dataset-specific patterns. CTSR mitigates this by cyclically transferring the shared backbone across related structural prediction tasks, including secondary structure and contact map prediction, before returning to the primary interface prediction objective. Each transfer introduces controlled perturbations to the optimization landscape while preserving shared geometric representations, allowing the model to escape suboptimal minima and integrate complementary structural signals. The resulting improvements in AUPR highlight the importance of optimization strategy, rather than model scale alone, in low-data structural learning.

Interpretability analyses provide additional insight into the representations learned by the MGA framework of VASCIF. The inferred amino acid interaction matrices reproduce established biochemical features of Ab–Ag interfaces, including enrichment of tyrosine- and serine-mediated contacts and broader polar complementarity across interacting partners. Predicted interface residue distributions closely match experimentally observed epitope compositions while diverging from background sequence frequencies. The emergence of these patterns without explicit physicochemical encoding suggests that the model internalizes genuine determinants of molecular recognition rather than superficial correlations.

Interestingly, antibody-only paratope prediction retains substantial predictive power, indicating that binding propensity is partially encoded within antibody variable domains themselves. Although antigen structure ultimately determines binding specificity and orientation, residue-level interaction propensity appears to arise largely from structural features intrinsic to antibody architecture. This has practical implications for early-stage antibody discovery workflows, where structural information for candidate antigens is often incomplete.

Beyond antibody–antigen modeling, the principles introduced here should extend to a broad class of imbalance-dominated learning problems. Dynamic Masking provides a general mechanism for adaptive feature sparsification that can be incorporated into graph neural networks, transformer-based language models, or convolutional architectures. Similar sparsity patterns arise in protein–ligand binding site prediction, post-translational modification localization, and nucleic acid–protein interface mapping, where functional residues constitute only a small subset of the structural landscape. In natural language processing, rare but semantically critical tokens may benefit from adaptive representation gating³¹. In computer vision, salient object regions often occupy only a limited portion of the image frame, suggesting that learnable masking mechanisms may enhance signal discrimination under extreme class imbalance³². The cyclic transfer strategy similarly generalizes to scenarios where labeled data are scarce but related auxiliary tasks are available, enabling controlled perturbations of the optimization landscape that improve generalization without increasing architectural complexity.

More broadly, this work contributes to the growing intersection between geometric deep learning and structural biology. Biomolecular systems are inherently structured objects defined by spatial relationships rather than linear sequences alone. Graph-based attention mechanisms and masked representation learning provide a natural framework for capturing these geometric dependencies while preserving physical interpretability. As structural datasets continue to expand through advances in cryo-electron microscopy, structural proteomics, and predictive modeling, approaches that combine geometric learning with biophysical insight will likely play an increasingly central role in understanding molecular recognition.

Some limitations remain. The current framework relies primarily on static crystallographic structures and does not explicitly account for conformational ensembles or binding-induced structural rearrangements. Energetic contributions such as solvent effects and entropic components are also not directly incorporated. Moreover, available Ab–Ag structural datasets remain modest compared with the scale of data used to train modern protein foundation models. Future integration of ensemble structural representations, energetic scoring functions, and glycan-aware modeling may further improve predictive accuracy and mechanistic insight.

In summary, this study reframes antibody–antigen interface prediction as a sparse structural learning problem and demonstrates that adaptive representation gating, cyclic optimization strategies, and principled capacity alignment can substantially improve predictive performance under extreme class imbalance. Beyond achieving state-of-the-art results, the framework reveals emergent biophysical interaction patterns, illustrating how sparsity-aware machine learning architectures can recover fundamental principles of molecular recognition directly from structural data. Improved residue-level interface prediction will accelerate antibody discovery and engineering by enabling rapid identification of candidate binding determinants from structural models, facilitating epitope mapping, affinity maturation strategies, and prioritization of therapeutic antibody designs. As structural datasets continue to expand, predictive frameworks that integrate physical insight with machine learning will increasingly support rational design in immunology and protein engineering.

## METHODS

### Data

For comparison with previous studies, we trained and evaluated our model on datasets used in prior work and additionally introduced a new dataset designed to reduce data leakage and improve generalization.

#### Paragraph-Expanded Dataset

The Paragraph dataset was derived from the Structural Antibody Database (SAbDab) using the March 31, 2022 release and contains 1,086 antibody-antigen (Ab–Ag) complexes^23^. The original training and validation sets were merged and evaluated using 5-fold cross-validation to address limited data availability. In the training–validation split, interface residues constitute 6.6% of all residues, with paratopes and epitopes comprising 5.1% and 10.6%, respectively. In the independent test set, these proportions are 6.4%, 5.1%, and 9.6%.

#### MIPE Dataset

The MIPE dataset was curated from SAbDab. Starting from 7,571 Ab–Ag complexes, redundancy was reduced using CD-HIT with 95% sequence identity for antibodies and 90% for antigens^25^. Complexes containing non-standard residues were removed, yielding 626 Ab–Ag pairs with sequence, structural, and interaction annotations. Residues were labeled as paratopes or epitopes if any backbone atom lay within 4.5 Å of a residue on the interacting partner. For model development, 90% of the data was used for 5-fold cross-validation and 10% reserved as an independent test set. Interface residues represent 6.1% of all residues in the training–validation set and 7.9% in the test set.

#### VASCO Dataset

To capture viral antigen diversity, we compiled an in-house dataset of antibody complexes against SARS, Influenza, Ebola, HIV, and Nipah antigens^28^. Because antibody sequences may share structural similarity, random splitting risks data leakage. We therefore embedded antibody sequences into vector representations, computed pairwise similarities, and clustered sequences using SciPy’s *fcluster*. Clusters were iteratively assigned to the training set until reaching 80% of the data, with the remainder used for testing. This cluster-based splitting reduces overfitting and improves generalization to unseen antigen structures.

### Input Representation

An Ab-Ag complex is represented as a sequence–structure pair (S,C), where S denotes the amino acid sequence and C the backbone coordinates. Binding residues are defined using a Euclidean distance threshold between antibody and antigen residues, with paratopes referring to antibody binding residues and epitopes to antigen binding residues. Given a dataset of complexes with sequence, structure, and annotated binding sites, the objective is to predict paratopes and epitopes for new Ab-Ag pairs. The model uses two input types: sequence matrices and adjacency matrices.

#### Sequence Matrices

Amino acid sequences from antibody and antigen chains are aligned using multiple sequence alignment (MSA). Alignments are generated with HHblits^45^ (three iterations, E-value cutoff 0.001). Homologous sequences are then clustered using k-means (Scikit-learn^46^) to maximize sequence diversity, following the strategy used in AlphaFold. Aligned sequences are represented as integer matrices of size (M×L), where M=64 is the number of homologous sequences used in this work and L=1600 is the maximum sequence length. Each row corresponds to one sequence in the MSA, and integers encode amino acids or special tokens (e.g., gaps or padding) following the ESM-Fold convention. Sequences are zero-padded to length L for batch processing. Matrices for the heavy chain, light chain, and antigen are stacked to form the full sequence representation (**Figure *8***).

#### Adjacency Matrices

Structural connectivity is represented as residue graphs. An edge is created between two residues if any pair of atoms lies within 10 Å. The resulting adjacency matrices are combined across chains and renumbered using end-of-chain symbols. Graph edges are stored in sparse format with shape [E,2] where E is the number of residue–residue connections.

### Target Representation

To improve model training, we use three prediction targets: secondary structure (SS), contact map (CTM), and interface residues (ITF). Secondary structures are assigned using STRIDE^47^, tokenized into eight classes, and padded to a fixed sequence length L=1600. The resulting label tensor has shape [L, 1]. The contact map captures all pairwise residue relationships, complementing adjacency matrices that encode only local connectivity. CTMs are represented as matrices of size [L, L], where each element denotes whether two residues are in contact. Interface residues are defined as residues with at least one heavy atom within 10 Å of any heavy atom on the interacting chain for comparison with previous studies. Interface labels are stored as vectors of size [L, 1]. These targets enable the model to learn complementary sequence and structural signals relevant to Ab–Ag interactions.

### Model Structure

The MGA architecture (**Figure 2**) employs dynamic masking and consists of an MSA module, a graph neural network (GNN) module, an attention module, and task-specific prediction layers, implemented as fully connected layers appended to the attention outputs.

### Embedding Layer

Input sequences are projected into a 256-dimensional embedding space with output shape [B, M, L, D], where B=4 is the batch size, M=64 the number of sequences, L=1600 the sequence length, and D=256 the embedding dimension.

### Dynamic Masking Architecture

Paratope and epitope prediction suffer from severe class imbalance, as only a small fraction of residues participates in binding. Conventional strategies such as restricting training to complementarity-determining regions (CDRs) or randomly down sampling non-binding residues rely on fixed heuristics and may limit generalization. To address this, we introduce a learnable Dynamic Masking (DyM) layer (**Figure 4**) integrated directly into the architecture. DyM enables the model to automatically identify informative residues while suppressing less relevant tokens in an end-to-end manner. Conceptually, the layer performs adaptive down-sampling, allowing the network to focus on residues most likely to contribute to binding. The approach is motivated by the observation that individual tokens carry intrinsic importance determined by structural and contextual features. By learning these patterns, DyM guides attention toward critical residues, improving both prediction accuracy and interpretability. The DyM module is implemented as a lightweight linear layer followed by a sigmoid mask applied to the embedding features.

### MSA Module

Embedding representations are processed through two consecutive self-attention operations within each MSA block: row-wise and column-wise attention. During row-wise attention, the input matrix is partitioned by chain so that attention is computed independently within each chain segment. The outputs are then recombined. From the resulting representation, the first row - corresponding to the embedding of the original sequence with shape [B, 1, L, D] - is extracted and passed to the next stage.

### GNN Module

The extracted sequence embedding is combined with a learnable positional embedding and used as input to the GNN module [B, 1, L, D]. This module applies multiple layers of Graph Attention Networks (GANs), where edges are defined by the adjacency matrices. The output retains the same dimensionality.

### Attention Module

An attention module composed of multiple self-attention layers further refines the representations. The final attention weights [B, H, L, L], where H=16 attention heads, are used for contact map prediction and visualization.

### Fully Connected Layers

Task-specific fully connected layers produce final predictions. For classification tasks (secondary structure or interface prediction), the output shape is [B, L, C], with C=8 or C=2. For contact map prediction, the input [B, H, L, L] is mapped to [B, 2, L, L].

### Model Training

All models were implemented in PyTorch and trained on four A40 GPUs. Training used the Paragraph, MIPE, and VISCO datasets with five-fold cross-validation at the complex level to prevent sequence overlap between folds. Optimization employed AdamW^48^ with a 10-epoch warm-up followed by cosine learning-rate decay. The batch size was 4, with gradient accumulation over two steps and gradient clipping at ∥g∥_2_ ≤ 0.5. To address class imbalance, a dynamic positive-class weighting scheme was applied. The loss function combined cross-entropy with label smoothing (ε=0.05), a differentiable AUPR surrogate (α = 0.4, k = 64 bins, temperature = 0.05), and a graph smoothness penalty encouraging similar predictions for neighboring residues. The smoothness coefficient (λ=0.05) was gradually introduced between epochs 5 to 15.

The overall loss function can be written as

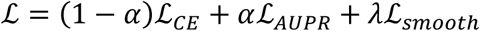

Here, ℒ_AUPR_ is differentiable surrogate for area under the PR curve. It is constructed by sweeping soft thresholds 𝑡_𝑘_ ∈ [0,1](with a sigmoid temperature) to get soft TP/FP/FN, then integrating precision over recall:

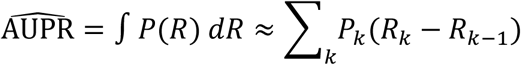

Where P and R are precision and recall in kth bin, and

ℒ_smooth_ is graph-based “region smoothing” penalty that encourages neighboring residues to have similar positive probabilities:

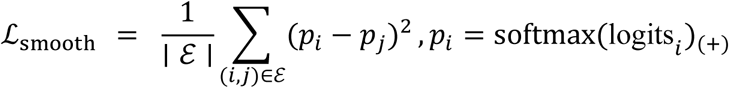

where ℰ are valid (unpadded) residue–residue edges.

To reduce overfitting and improve robustness, we applied graph-specific regularization, including random edge dropout (p = 0.05 – 0.15) and token masking (p = 0.1) during training, excluding the query MSA row. Stochastic Weight Averaging (SWA) and an exponential moving average (EMA) of model parameters (decay = 0.995) were maintained throughout training. Final predictions were obtained by averaging EMA and SWA outputs to improve calibration. Model selection within each fold was based on the highest validation AUPR.

To maximize performance under limited labeled data, we introduce Cyclic Task Self-Refinement (CTSR). The model is first trained on the primary task of interface (ITF) prediction, and the learned backbone parameters (excluding the task-specific layer) are transferred to an auxiliary task such as secondary structure (SS) prediction. After auxiliary training, the updated backbone is transferred back to the original task, and the process can continue with additional tasks such as contact map (CTM) prediction. CTSR is motivated by the observation that small datasets can cause gradient descent to become trapped in local minima^49^. Related tasks can be viewed as slightly perturbed versions of the same optimization landscape. Transferring between tasks effectively shifts this landscape, enabling the model to escape suboptimal minima while integrating complementary structural signals. This cyclic transfer improves generalization while maximizing the value of limited structural data.

### Training Procedure

#### 1. Initial ITF Training

For residue-level interface prediction we used LR = 2×10⁻⁴, 50 epochs maximum, SWA starting at epoch 20. The positive weight decayed exponentially from 20 to 1 with a time constant τ = 8.

#### 2. Pretraining for Secondary Structure Prediction (SS)

We initialized from the best ITF checkpoints (all weights except the final classification layer) and fine-tuned on SS with cross-entropy, using a reduced learning rate (1×10⁻⁴) and the same regularization stack (dropout, drop-path, gradient noise, clip). For 8-class secondary-structure prediction we used: batch size = 4, LR = 1×10⁻⁴, 50 epochs maximum, SWA from epoch 20. Class weights were fixed to 1.0 (no annealing).

#### 3. Iterative Training on Multiple Targets

Starting from the SS-tuned backbone, we returned to ITF for a short fine-tune with reduced LR and increased dropout and drop-path rate to preserve and sharpen interface discrimination, then trained CTM, and finally performed one additional ITF fine-tune. This loop allows the backbone to absorb complementary structural signals (secondary structure and long-range contacts) while preventing drift of the interface head.

## Funding and Acknowledgements

X.L., J.S. and S.C. were supported by NIH NIGMS grant R35GM151231-01, and through NEU Faculty Startup Funds. This research used computational resources from Northeastern Discovery cluster at MGHPCC, as well as through NSF ACCESS.

## Author contributions

X.L. A.K.C. and S.C. conceptualized the study and designed the experiments. X.L. prepared and tested the ML model. X.L. and J.K performed data analysis. X.L. and S.C. performed data interpretation. J.K. and X.L. prepared the figures. X.L. and S.C drafted the manuscript. X.L., J.K., A.K.C and S.C. read and edited the final manuscript. S.C. is the corresponding author.

## Data and materials availability

All codes and materials are shared freely on github: https://github.com/SimBioSys-lab/MGA

A webserver version for prediction can also be found at: https://glacier-simbiosys.com/vasco

Further details as required can be requested from the authors.

