## Supplemental Table 1 for "Interpretable Antibody–Antigen Structural Interface Prediction via Adaptive Graph Learning and Cyclic Transfer"

### Supplementary material

**Table S1 Performance of models with various structure on three datasets**

| Dataset | MSA layers | GNN layers | Attention layers | Antibody AUPR | Antibody AUC-ROC | Antigen AUPR | Antigen AUC-ROC |
|---|---|---|---|---|---|---|---|
| MIPE | 1 | 4 | 2 | 0.725 | 0.981 | 0.247 | 0.753 |
| | 1 | 6 | 2 | 0.732 | 0.981 | 0.256 | 0.758 |
| | 1 | 6 | 3 | 0.742 | 0.981 | 0.282 | 0.775 |
| | 1 | 6 | 4 | 0.726 | 0.981 | 0.222 | 0.757 |
| | 1 | 8 | 2 | 0.729 | 0.981 | 0.244 | 0.759 |
| | 1 | 8 | 3 | 0.722 | 0.981 | 0.246 | 0.761 |
| | 1 | 8 | 4 | 0.726 | 0.981 | 0.223 | 0.757 |
| | 1 | 10 | 5 | 0.726 | 0.981 | 0.215 | 0.747 |
| paragraph | 0 | 10 | 5 | 0.747 | 0.982 | 0.449 | 0.801 |
| | 1 | 0 | 5 | 0.584 | 0.964 | 0.409 | 0.766 |
| | 1 | 10 | 0 | 0.726 | 0.980 | 0.454 | 0.798 |
| | 1 | 10 | 5 | 0.751 | 0.982 | 0.472 | 0.812 |
| | 1 | 2 | 1 | 0.721 | 0.980 | 0.461 | 0.810 |
| | 1 | 4 | 2 | 0.735 | 0.981 | 0.458 | 0.806 |
| | 1 | 6 | 3 | 0.747 | 0.982 | 0.461 | 0.809 |
| | 1 | 8 | 4 | 0.746 | 0.982 | 0.458 | 0.803 |
| | 1 | 10 | 4 | 0.752 | 0.982 | 0.469 | 0.810 |
| | 1 | 10 | 6 | 0.753 | 0.982 | 0.464 | 0.811 |
| | 1 | 12 | 4 | 0.752 | 0.982 | 0.452 | 0.798 |
| | 1 | 12 | 5 | 0.752 | 0.982 | 0.458 | 0.801 |
| | 1 | 12 | 6 | 0.753 | 0.982 | 0.461 | 0.805 |
| | 1 | 14 | 7 | 0.753 | 0.982 | 0.452 | 0.799 |
| VASCO | 1 | 10 | 5 | 0.716 | 0.795 | 0.528 | 0.943 |
| | 1 | 12 | 6 | 0.730 | 0.964 | 0.541 | 0.932 |
| | 1 | 14 | 7 | 0.700 | 0.963 | 0.498 | 0.936 |
| | 1 | 16 | 8 | 0.693 | 0.963 | 0.464 | 0.931 |
